# Neural oscillations as a signature of efficient coding in the presence of synaptic delays

**DOI:** 10.1101/034736

**Authors:** Matthew Chalk, Boris Gutkin, Sophie Deneve

## Abstract

Cortical networks exhibit “global oscillations”, in which neural spike times are entrained to an underlying oscillatory rhythm, but where individual neurons fire irregularly, on only a fraction of cycles. While the network dynamics underlying global oscillations have been well characterised, their function is debated. Here, we show that such global oscillations are a direct consequence of optimal efficient coding in spiking networks with synaptic delays. To avoid firing unnecessary spikes, neurons need to share information about the network state. Ideally, membrane potentials should be strongly correlated and reflect a “prediction error” while the spikes themselves are uncorrelated and occur rarely. We show that the most efficient representation is achieved when: (i) spike times are entrained to a global Gamma rhythm (implying a consistent representation of the error); but (ii) few neurons fire on each cycle (implying high efficiency), while (iii) excitation and inhibition are tightly balanced. This suggests that cortical networks exhibiting such dynamics are tuned to achieve a maximally efficient population code.

## Introduction

Oscillations are a prominent feature of cortical activity. In sensory areas, one typically observes “global oscillations” in the gamma-band range (30-80Hz), alongside single neuron responses that are irregular and sparse [1, 2]. The magnitude and frequency of gamma-band oscillations are modulated by changes to the sensory environment (e.g. visual stimulus contrast [3]) and behavioural state (e.g. attention [4]) of the animal. This has led a number of authors to propose that neural oscillations play a fundamental role in cortical computation [5, 6]. Others argue that oscillations emerge as a consequence of interactions between populations of inhibitory and excitatory neurons, and do not perform a direct functional role in themselves [7].

A prevalent theory of sensory processing, the “efficient coding hypothesis”, posits that the role of early sensory processing is to communicate information about the environment using a minimal number of spikes [8]. This implies that the responses of individual neurons should be as asynchronous as possible, so that they do not communicate redundant information [9]. Thus, oscillations are generally seen as a bad thing for efficient rate coding, as they tend to synchronise neural responses, and thus, introduce redundancy.

Here we propose that global oscillations are a necessary consequence of efficient rate coding in recurrent neural networks with synaptic delays.

In general, to avoid communicating redundant information, neurons driven by common inputs should actively decorrelate their spike trains. To illustrate this, consider a simple set-up in which neurons encode a common sensory variable through their firing rates, with a constant value added to the sensory reconstruction each time a neuron fires a spike (**Fig. 1a**).

**Fig. 1.**
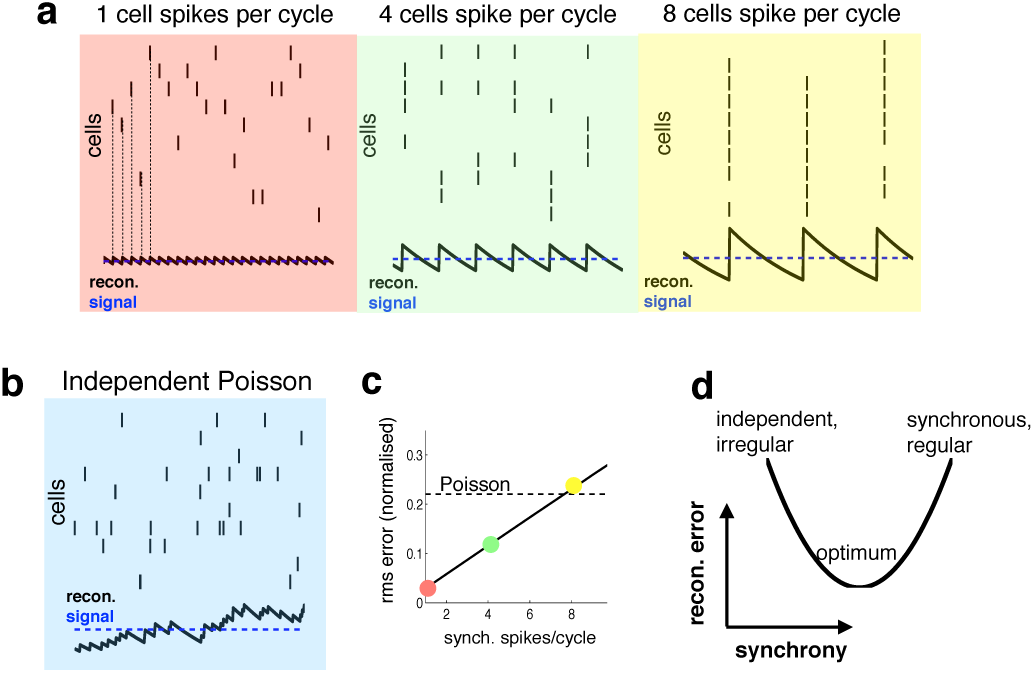
Relationship between synchrony and coding accuracy, (a) Each panel illustrates the response of 10 neurons, An encoded sensory variable is denoted by a horizontal blue line, Each spike fired by the network increases the sensory reconstruction by a fixed amount, before it decays, Greatest accuracy is achieved when the population fires at regular intervals, but no two neurons fire together (left), Coding accuracy is reduced when multiple neurons fire together (middle and right panels), (c) Same as panel a, but where neurons show independent poisson variability, (b) Reconstruction error (root-mean squared error divided by the mean) for a regular spiking network shown in panel a, versus the number of synchronous spikes on each cycle, A horizontal line denotes the performance when neurons fire with independent Poisson variability, (c) Cartoon illustrating tradeoff between trade-off faced by neural networks.

With a constant input, the reconstruction error is minimized when the population fires spikes at regular intervals, while no two neurons fire spikes at the same time (as in **Fig 1a**, left panel, in red). To achieve this ideal, however, requires incredibly fast inhibitory connections, so that each time a neuron fires a spike it suppresses all other neurons from firing [10]. In reality, inhibitory connections are subject to unavoidable delays (e.g. synaptic and transmission delays), and thus, cannot always prevent neurons from firing together. Worse, in the presence of delays, inhibitory signals, intended to prevent neurons from firing together, can actually have the reverse effect of *synchronising* the network, so that many neurons fire together on each cycle (as in **Fig. 1a**, middle and right panels). This is analogous to the so-called ‘hipster effect’ where a group of individuals strive to look different from each other, but due to delayed reaction times, end up making similar decisions and all looking alike [11].

Spiking synchronicity generally has a negative effect on coding performance. For example, **figure 1c** shows how, in the regular spiking network described above, coding error increases with the number of synchronous spikes per cycle (while firing rate is held constant). It is thus tempting to conclude that neural networks should do everything possible to avoid synchronous firing. However, one also observes that a completely asynchronous network, in which neurons fire with independent Poisson variability (**Fig. 1b**), performs far worse than the regular spiking network, even when multiple neurons fire together on each cycle (**Fig. 1c**, horizontal dashed line).

Thus, to perform efficiently, neural networks face a tradeoff (**Fig. 1d**). On the one hand recurrent connections should coordinate the activity of different neurons in the network, so as to achieve an efficient and accurate population code. On the other hand, in the presence of synaptic delays, it is important that these recurrent signals do not overly synchronize the network, as this will reduce coding performance.

Here, we show that, in a network of leaky integrate-and-fire neurons (LIF) optimized for efficient coding, this trade-off is best met when: (i) neural spike trains are entrained to a global oscillatory rhythm (for a consistent representation of global information), but (ii) only a small fraction of cells fire on each oscillation cycle. In this regime, individual neurons fire irregularly [12], and exhibit weak pairwise correlations [13], despite the presence of rhythmic population activity. Moreover, excitation and inhibition are tightly balanced on each oscillation cycle, with inhibition lagging excitation by a few milliseconds [14, 15]. Thus, ‘global oscillations’ come about as a direct consequence of efficient rate coding, in a recurrent network with synaptic delays.

## Results

**Efficient coding in an idealized recurrent network**. It is instructive to first consider the behaviour of an idealized network, with instaneous synapses. For this, we consider a model proposed by Boerlin et al., in which a network of integrate-and-fire neurons is optimized to efficiently encode a time varying input. As the model has already been described elsewhere [10], we restrict ourselves to outlining the basic principles, with a mathematical description reserved for the **Methods**.

Underlying the model is the idea that downstream neurons should be able to reconstruct the input to the network by performing a linear summation of its output spike trains. To do this efficiently (i.e. with as few spikes as possible), the spiking output of the network is fed-back and subtracted from its original input (**Fig. 1a**). In consequence, the total input to each neuron is equal to a ‘prediction error’; the difference between the original input and the network reconstruction. This prediction error is also reflected in neural membrane potentials. When a neuron’s membrane potential becomes larger than a constant threshold then it fires a spike; recurrent feedback then reduces the prediction error encoded by other neurons, preventing them from firing further redundant spikes.

To illustrate the principles underlying the model, we consider a network of three identical neurons. **Figure 2b** shows how spikes fired by the network are ‘read-out’, to obtain a continuous reconstruction. Each time a neuron fires a spike, it increases the reconstruction by a fixed amount, decreasing the difference between the input and the neural reconstruction. Immediately after, feedback connections alter the membrane potentials of other neurons, to ensure that they maintain a consistent representation of the error, and do not fire further spikes (**Fig. 2b**, lower panel). As a result, membrane potentials are highly correlated, while spikes are asynchronous.

**Fig. 2.**
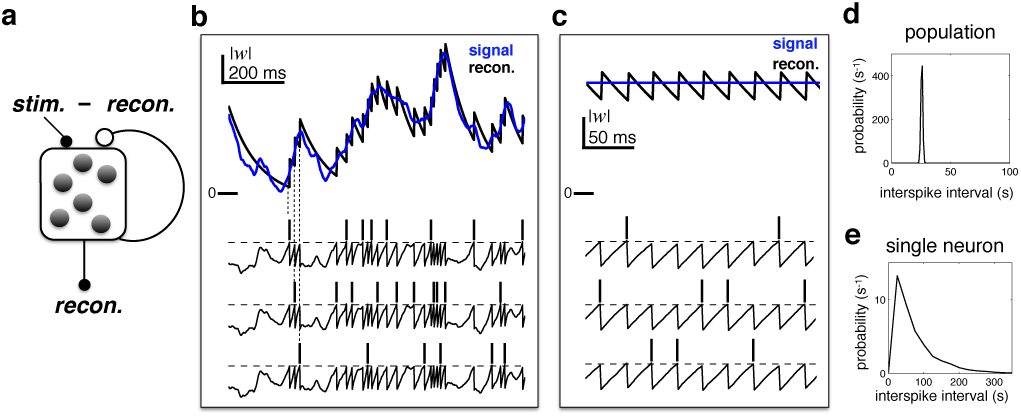
Efficient coding in a recurrent neural network. (**a**) Schematic of network. Inhibitory recurrent connections are represented by an open circle. Excitatory feed-forward connections are represented by closed circles. (**b**) Stimulus (blue) and neural reconstruction (black) on a single trial. The spikes and membrane potential for each each cell are shown in seperate rows. Vertical dashed lines illustrate how each spike alters the neural reconstruction. (**c**) Same as (**b**), but with a constant input (also note the change of temporal scale). (**d**-**e**) Distribution of inter-spike intervals in population (**d**) and single-cell (**e**) spike trains.

**Figure 2c** shows the behaviour of the network in response to a constant input. To optimally encode a constant input, the network generates a regular train of spikes (as in the left panel of **Fig. 1a**), resulting in a narrow distribution of population inter-spike intervals (ISIs) (**Fig. 2d**). Neural membrane potentials, which encode a common prediction error, fluctuate in synchrony, with a frequency dictated by the population firing rate (**Fig. 2c**, lower panel). However, as only one neuron fires per cycle, the spike trains of *individual neurons* are irregular and sparse, resulting in a near-exponential distribution of single-cell ISIs (**Fig. 2e**).

**Efficient coding with synaptic delays**. In real neural networks, recurrent inhibition is not instantaneous, but subject to synaptic and transmission delays. Far from being a biological detail, even very short synaptic delays can profoundly change the behaviour of the idealized efficient coding network, and pose fundamental limits on its performance.

To render the idealized model biologically feasible we extended it in two ways. First, to comply with Dale’s law, we introduced a population of inhibitory neurons, which mediates recurrent inhibition (**Fig. 3a**). In our implementation, excitatory and inhibitory populations both encode separate reconstructions of the target variable. Inhibitory neurons, which receive input from excitatory neurons, fire spikes in order to predict and cancel the inputs to the excitatory population (see **Methods**).

**Fig. 3.**
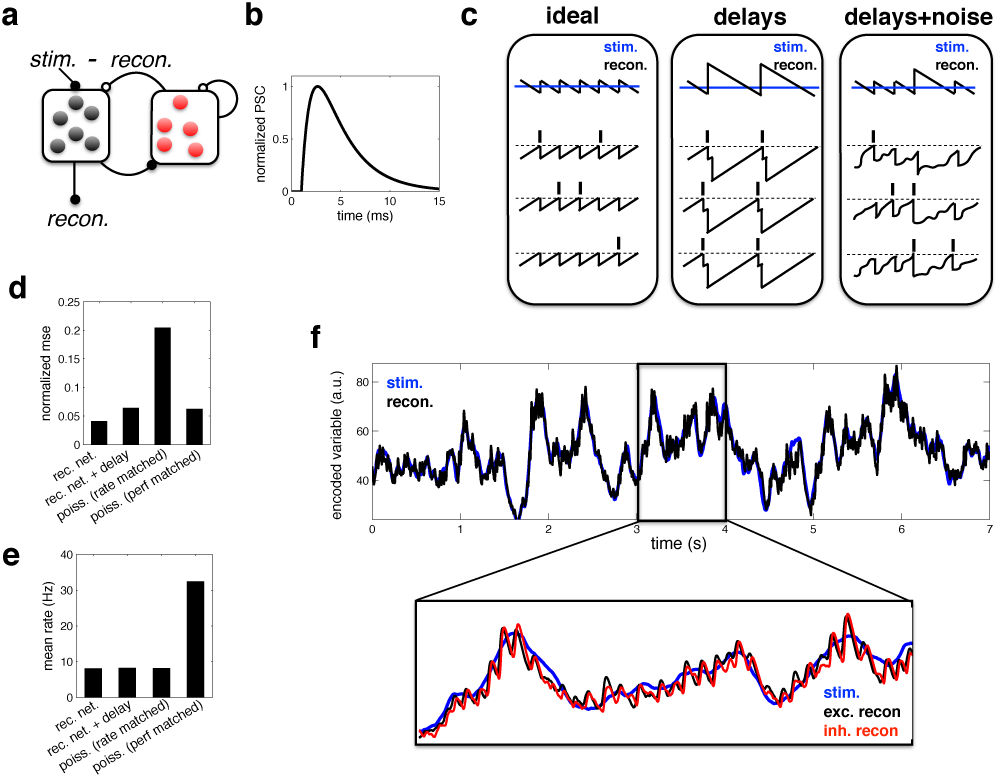
Coding performance of an excitatory/inhibitory network with synaptic delays. (**a**) Schematic of network connectivity. Excitatory neurons and inhibitory neurons are shown in black and red, respectively. Connections between different neural populations are schematised by lines terminating with solid (excitatory connections) and open (inhibitory connections) circles. (**b**) The postsynaptic current (PSC) waveform used in our simulations. (**c**) Schematic, illustrating how delays affect the decoding performance of the network. (**d**) Normalised mean squared reconstruction error and (**e**) population firing rate for (i) an ideal network with instantaneous synapses, (ii) a network with finite synaptic delays, (iii) independent Poisson units whose firing rate varies as a fixed function of the feed-forward input. We compare two types of Poisson model, one with average firing rates that match the recurrent network (‘rate matched’), and one with firing rates scaled to match the encoding performance of the recurrent model (‘performance matched’). (**f**) Stimulus and neural reconstruction for the efficient coding network, with synaptic delays. (**inset**) Excitatory (black) and inhibitory (red) neural reconstructions, during a short segment from this trial.

Second, and more importantly, we replaced the instantaneous synapses in the idealized model with continuous synaptic dynamics, as shown in **figure 3b** (see **Methods**). As stated previously, adding synaptic delays substantially alters the performance of the network. Without delays, recurrent inhibition prevents all but one cell from firing per oscillation cycle, resulting in an optimally efficient code (**Fig. 3c**, left panel). With delays however, inhibition cannot always act fast enough to prevent neurons firing together. As a result, neural activity quickly becomes synchronised, and the sensory reconstruction is destroyed by large population-wide oscillations (**Fig. 3c**, middle panel). To improve performance, one can increase the membrane potential noise and spike threshold, so as to reduce the chance of neurons firing together (**Fig. 3c**, right panel). Too much noise, however, and the firing of different neurons becomes uncoordinated, and network performance is diminished (see later).

We compared the performance of the efficient coding network (with excitatory/inhibitory populations and synaptic delays) to: (i) an ‘ideal’ model with no delays and (ii) a ‘rate model’, consisting of a population of independent Poisson units whose firing rate varies as a function of the feed-forward input (see **Methods**). **Figure 3d-e** show the average firing rate and reconstruction error achieved with each model type, in response to a time varying input. Finite synaptic delays moderately increased the reconstruction error (**Fig. 3d**) but did not substantially change the average firing rate (**Fig. 3e**). Despite this increase in coding error, the recurrent network still performed significantly better than a network of independent Poisson units, with matched rates (‘rate matched’ model) or reconstruction error (‘performance matched’ model).

**Figure 3f** illustrates the ability of the network to track a time varying input signal. Zooming in to a 1s period within the trial (lower inset), one observes rhythmic fluctuations in the excitatory and inhibitory neural reconstructions. These fluctuations are essentially the same phenomena as observed for the ideal network, where the neural reconstruction fluctuated periodically around the target signal following the arrival of each new spike (**Fig. 2**). However, with synaptic delays, several neurons fire together before the arrival of recurrent inhibition. As a result, oscillations are slower and larger in magnitude than for the idealised network, where only one neuron fires on each cycle.

**Oscillations and efficient coding**. We sought to quantify the effect of oscillations on coding performance. To do this, we varied parameters of the model, so as to alter the degree of network synchrony, while keeping firing rates the same. In the main text we illustrate the effect of adding white noise to the membrane potentials (while simultaneously varying the spike threshold, to maintain constant firing rate; see **Methods**). Qualitatively similar results were obtained by altering other aspects of the network, such as varying the probability of synaptic failure (**Supp. fig. 2**) or the ratio of feed-forward to recurrent inhibition (**Supp. fig. 3–4**). Further, while for simplicity we considered the response to a constant input in the main text, qualitatively similar results were obtained with a time-varying input (see, for example, **Supp. fig. 6**).

Increasing the magnitude of the membrane potential noise desynchronized the network activity, resulting in a reduction in pairwise voltage correlations (**Supp fig. 1a**). With increased noise, single neural spike trains also became more irregular, reflected by an increase in the spiking CV (**Supp fig. 1b**).

Coding performance, however, varied non-monotonically with the noise amplitude. The reconstruction error followed a u-shaped curve, being minimised for a certain intermediate level of noise (**Fig. 4a**, solid curve). For this intermediate noise level, coding performance was significantly better than a network of independent Poisson units with matching firing rate (horizontal dashed line). Interestingly, deviations between excitation and inhibition followed a similar u-shape curve, being minimised for the same intermediate noise level (**Fig. 4a**, thick dashed curve). Thus, optimal coding was achieved when the balance between excitatory and inhibition was the tightest.

**Fig. 4.**
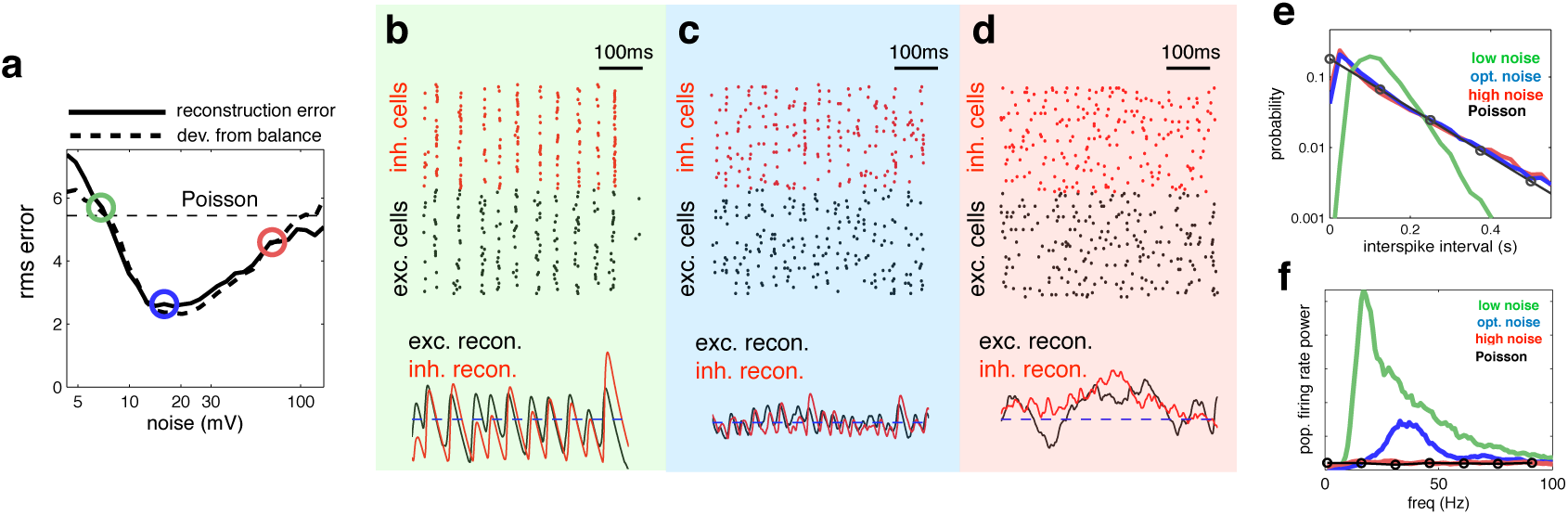
Effect of varying noise amplitude on network dynamics and coding performance. (**a**) Root-mean-square (rms) reconstruction error (solid line), plotted alongside rms difference between excitatory and inhibitory reconstructions (dashed line), versus injected noise amplitude. Horizontal dashed line shows the rms reconstruction error for a population of Poisson units, with identical firing rates. (**b**-**d**) (above) Spiking response of inhibitory (red) and excitatory (black) neurons. (below) Inhibitory (red) and excitatory (black) neural reconstructions, alongside target stimulus (blue dashed line). Each plot corresponds to a different noise level, indicated by the coloured circles in panel **a**. (**e**) Distribution of single-cell inter-spike intervals in each noise condition. The prediction for a population of Poisson units is shown in black. Note the log-scale. (**f**) Power spectrum of population firing rate, in each noise condition. The power spectrum for a population of Poisson units is shown in black.

We quantified the degree to which population and single neuron responses differed from Poisson statistics (**Supp fig 1c**). Interestingly, we found that optimal coding performance (indicated by blue circles) was achieved when individual neuron spike trains were effectively Poisson, but where the population response was non-Poisson. This reflects the fact that, in the optimal regime, the population response exhibited global oscillations (and was thus, highly non-Poisson), while single-cell responses were irregular and sparse (and thus, close to Poisson).

To understand the effect of varying noise amplitude, we plotted the network responses and neural reconstruction in three regimes: with low, intermediate, and high noise (indicated by green, blue and red circles respectively in **Fig. 4a**).

With low noise, neural membrane potentials were highly correlated, leading many neurons to fire together on each oscillation cycle (**Fig 4b**, upper panel). As a result, the neural reconstruction exhibited large periodic fluctuations about the encoded input, leading to poor coding performance (**Fig**. **4b**, lower panel).

On the other extreme, when the injected noise was very high, the spike trains of different neurons were uncorrelated (**Fig 4d**, upper panel). As, in this regime, effectively no information was shared between neurons, inhibitory and excitatory reconstructions were decoupled, and coding performance was similar to a population of independent Poisson units (**Fig 4d**, lower panel).

In the intermediate noise regime, for which performance was optimal, spikes were aligned to rhythmic fluctuations in the prediction error, but few neurons fired on each cycle (**Fig 4c**, upper panel). These dynamics were reflected by a near-exponential distribution of interspike-intervals (**Fig. 4f**), coupled with a narrow peak in the population firing rate spectrum (**Fig 4e**). In this regime, rhythmic fluctuations in the neural reconstruction were small in magnitude, and there was a tight coupling between inhibitory and excitatory reconstructions (**Fig**. **4c**, lower panel).

**Oscillatory neural dynamics**. We investigated the behaviour of the network in the optimal regime, shown in figure 4c. **Figure 5a** shows the network reconstruction and population firing rates in response to a low (blue), medium (red) and high (green) amplitude stimulus. The population firing rate was characterised by a transient peak following stimulus onset, followed by decay to a constant value. A spectrogram of the population firing rate (**Fig. 5a**, lower panel) reveals the presence of 30-50Hz oscillations during the period of sustained activity. **Figure 5b** plots the spiking response and population firing rates during a 600ms period of sustained activity. Here, one clearly sees correlated rhythmic fluctuations in excitatory (black) and inhibitory (red) activity. The strength of these oscillations increases with stimulus amplitude (**Fig. 5c**). Nevertheless, for all input amplitudes, individual neurons fired irregularly, with a near-exponential distribution of inter-spike intervals (**Fig. 5d**).

**Fig. 5.**
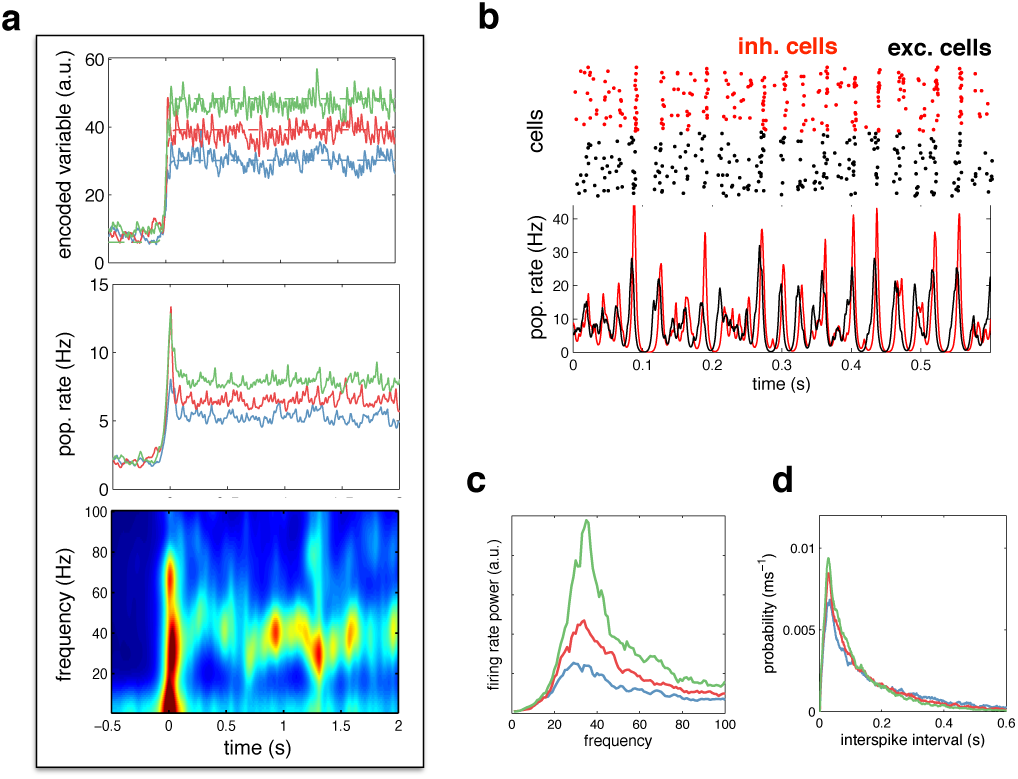
Spiking response to a constant input, (**a**) (**top**) Neural reconstruction (solid lines) of a presented constant stimulus (dashed lines), before and after stimulus onset, Low, medium and high amplitude stimuli are shown in blue, red and green, respectively, (**middle**) Population firing rate, for each stimulus amplitude, (**bottom**) Spectrogram of population firing rate, (**b**) The upper panel shows a raster-gram of excitatory (black) and inhibitory (red) responses, during a 0,6s period of sustained activity. The lower panel shows the instantaneous firing rates of the excitatory and inhibitory populations, (**c**) Power spectrum of excitatory population firing rate during the sustained activity period, for each stimulus amplitude, (**d**) Average distribution of single-cell inter-spike intervals, for each stimulus amplitude.

We next considered the dynamics of neural membrane potentials. Previously, Yu & Ferster [2] reported that, in area V1, visual stimulation increases gamma-band correlations between pairs of neural membrane potentials (**Fig**. **6a**). Qualitatively similar results were obtained with our model (**Fig. 6b**). Increasing the amplitude of the feed-forward input led to increased correlations between neural membrane potentials (**Fig. 6c**), with strongest coherence observed in the gamma-band range (**Fig. 6d**). This is because more neurons fire spikes on each cycle, leading to stronger oscillations.

**Fig. 6.**
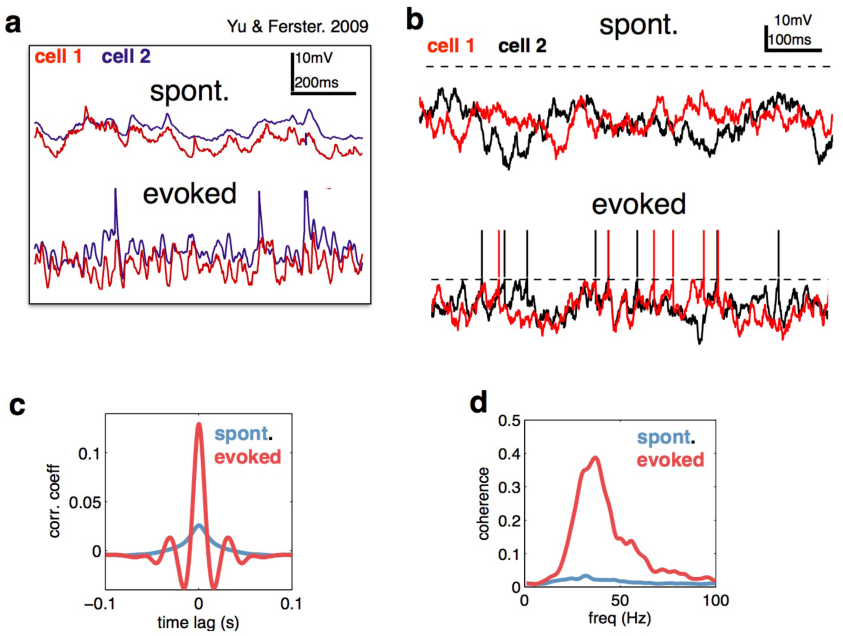
Voltage responses to a constant input, (**a**) Data from Yu & Ferster (Neuron, 2010), showing voltage traces of two V1 cells, in the absence (top), and presence (bottom) of a visual stimulus, (**b**) Voltage traces of two cells in the model in absence (top) and presence (bottom) of a feed-forward input, (**c**) Average pairwise cross-correlation between membrane potentials, in spontaneous and evoked condition, (**e**) Average coherence spectrum between pairs of voltage traces, in spontaneous and evoked condition.

Several studies have reported a tight balance between inhibition and excitation [14, 15, 16]. Recently, Atallah et al. [14] reported that inhibitory and excitatory currents are precisely balanced on individual cycles of an ongoing gamma oscillation (**Fig. 7a**). In our model, efficient coding is achieved by maintaining such a tight balance between inhibitory and excitatory reconstructions. Thus, inhibitory and excitatory currents closely track each other (**Fig. 7b**), with a high correlation between the amplitude of inhibitory and excitatory currents on each cycle (**Fig. 7c**). In common with Atallah et al.’s data, inhibition lags behind excitation by a few milliseconds (**Fig. 7d**). Fluctuations in the amplitude of inhibitory and excitatory currents instantaneously modulate the oscillation frequency, with a significant correlation observed between the peak amplitude on a given oscillation cycle and the period of the following cycle (**Fig. 7e**).

**Fig. 7.**
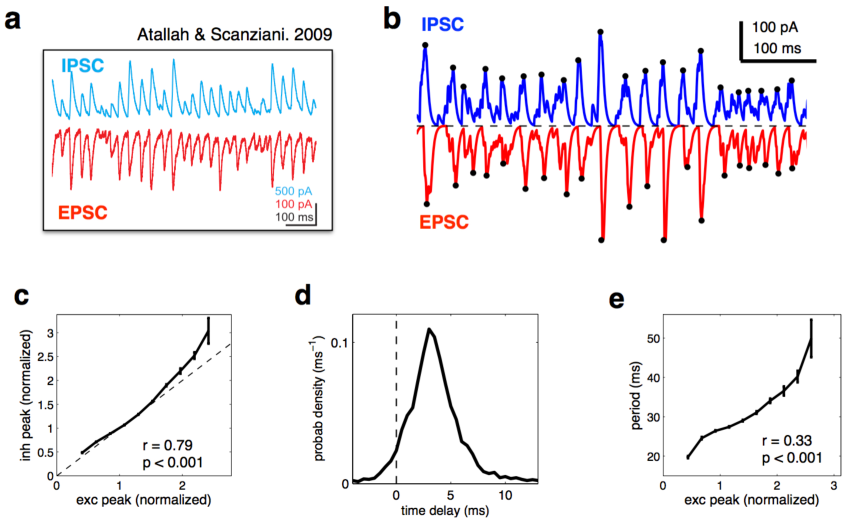
Balanced fluctuations in excitatory and inhibitory currents, (**a**) Data from Atallah & Scanziani (Neuron, 2009), showing inhibitory (blue) and excitatory (red) postsynaptic currents, (**b**) Inhibitory (blue) and excitatory currents (red) in the model, in response to a constant feed-forward input, Black dots indicate detected peaks, (**b**) Distribution of periods between successive peaks in excitation, (**c**) Amplitude of inhibitory current versus magnitude of excitatory currents on each cycle, (**d**) Distribution of time lags between excitatory and inhibitory peaks, (**e**) Period between excitatory peaks, versus the peak amplitude on the previous oscillation cycle.

**Gamma oscillations and behavioural performance**. In general, the optimal network parameters depend on the properties of the feed-forward sensory input. For example, the higher the input amplitude, the more noise is required to achieve the optimal level of network synchrony (**Fig. 8a**). While the network achieves reasonable coding accuracy for a large range of different inputs, adaptive tuning of the dynamics (for example, changing the noise level) can be beneficial for a more limited input range. This would affect the level of population synchrony and thus introduce a correlation between performance and the strength of Gamma oscillations.

**Fig. 8.**
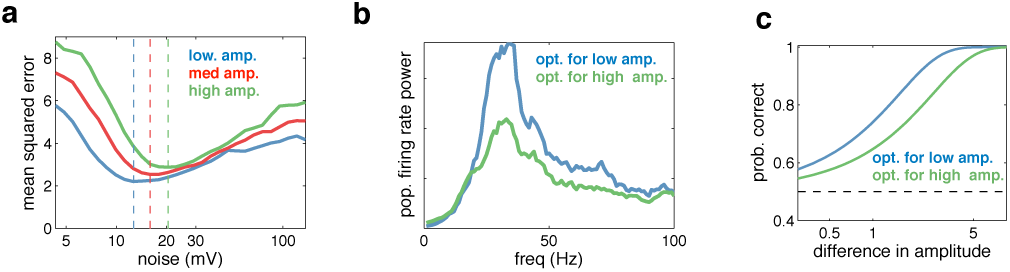
Effect of varying the input amplitude. (**a**) Root-mean-square (rms) reconstruction error versus injected noise amplitude with three different input amplitudes. The optimal noise level in each condition is denoted by vertical dashed lines. (**b**) Power spectrum of population firing rate, in response to a low amplitude input. The blue curve corresponds to when the network is optimised for the low amplitude input, the green curve for when it is optimised for the high amplitude input. (**c**) Fractional performance in discriminating between two 100ms input segments, equally spaced around the low amplitude input. Performance is best in the condition with strongest oscillations.

For example, if the task is to detect weak (low amplitude) inputs, performance would be higher if top down modulations (such as attention) reduced the level of noise, and thus increased the degree of population synchrony (**Fig. 8b**) without significantly changing firing rates. We can thus expect higher detection performance to correlate with stronger level of Gamma oscillations (**Fig 8c**). This could account for certain attention-dependent increases in gamma-power and its correlation with behavioural performance (see **Discussion**).

Note that a similar correlation between behavioural performance and Gamma power could arise from purely bottom up effects. In the presence of input noise causing trial-by-trial changes in input strength, trials with stronger input amplitude would result in more detection but also exhibit more Gamma oscillations. In that case, however, increase in Gamma power would be associated with a commensurate increase in population firing rate.

## Discussion

We present a novel hypothesis for the role of neural oscillations, as a consequence of efficient coding in recurrent neural networks with noise and synaptic delays. In order to efficiently code incoming information, neural networks must trade-off two competing demands. On the one hand, to ensure that each spike conveys new information, neurons should actively desynchronise their spike trains. On the other hand, to do this optimally, neural membrane potentials should encode shared global information about what has already been coded by the network, which will tend to synchronise neural activity.

In a network of LIF neurons with dynamics and connectivity tuned for efficient coding, we found that this trade-off is best met when neural spike trains are entrained to a global oscillatory rhythm (implying a consistent representation of the prediction error), but where few neurons fire spikes on each cycle (implying high efficiency). This also corresponds to the regime in which inhibition and excitation are most tightly balanced. Our results provide a functional explanation for why cortical networks operate in a regime in which: (i) global oscillations in population firing rates occur alongside individual neurons with low, irregular, firing rates [17] (ii) there is a tight balance between excitation and inhibition [14, 15, 16].

For simplicity, we considered a homogeneous network with one-dimensional feed-forward input. However, the results presented here easily generalise to networks with heterogenous connection strengths (**Supp. fig 5**), as well as networks that encode high-dimensional dynamical variables [10] (**Supp. fig 6**).

**Relation to balanced network models**. Previously, Brunel & colleagues derived the conditions under which a recurrent network of integrate-and-fire neurons with sparse irregular firing rates exhibits fast global oscillations [17, 18, 19, 20]. This behaviour is qualitatively similar to the network dynamics observed in our model. However, these previous models differ in several ways from the model presented here. For example, they assume sparse connections (and/or weak connectivity), in which the probability of connections (and/or connection strengths) scales inversely with the number of neurons. In contrast, the connections in our network are non-sparse and finite. Thus, our network achieves a tighter balance between inhibitory and excitatory currents, and smaller fluctuations in membrane potentials (they scale as 1/*N* in the absence of delays, rather than 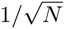).

However, the most important distinction between our work and previous mean-field models lies in the way the network is constructed. In our work, the network connectivity and dynamics are derived from functional principles, in order to minimise a specific loss function (i.e. the squared difference between the neural reconstruction and input signal). This ‘top-down’ modeling approach allows us to directly ask questions about the network dynamics that subserve optimal efficient coding. For example, balanced inhibition and excitation are not imposed in our model, but rather, required for efficient coding. Further, while previous models showed mechanistically how fast oscillations can emerge in a network with slow irregular firing rates [17], our work goes further, showing that these dynamics are in fact required for optimal efficient coding.

Finally, it is important to realise that, while efficient coding in a recurrent network leads to global oscillations, the reverse is not true: just because a network oscillates, does not mean that it is performing efficiently. To demonstrate this point, we repeated our simulations in a network with heterogeneous read-out weights (**Supp. fig 5**). Both the coding performance and spiking dynamics of this network were indistinguishable from the homogeneous network described in the main text. In contrast, when we randomised the recurrent connection strengths (while keeping the total input to each neuron the same), the coding performance of the network was greatly reduced, despite the fact that the network dynamics and firing rate power spectrum were virtually unchanged.

The coherence between excitatory and inhibitory current oscillations (i.e. the level of balance) is a much more reliable signature of efficient coding than global population synchrony. While population synchrony can occur in globally balanced network as well, only networks with intracellular detailed balance between excitatory and inhibitory currents achieve high coding performance (Supp. fig. 5b & d).

**Relation to previous efficient coding models**. Previous work on efficient coding has mostly concentrated on using information theory to ask what information ‘should’ be represented by sensory systems [9]. Recently, however, researchers have begun to ask, mechanistically, how neural networks should be setup in order to operate as efficiently as possible [10, 21, 22]. This approach can provide certain insights not obtainable from information theory alone.

For example, information theory suggests that the most efficient spiking representation is one in which the spike trains of different neurons are statistically independent, and thus, there are no oscillations. In practice however, neural networks must operate in the face of biological constraints, such as synaptic delays and noise. Considering these constraints changes our conclusions. Specifically, contrary to what one would expect from a purely information theoretical analysis, we find that oscillations emerge as a consequence of neurons performing as efficiently as possible given synaptic delays, and should not be removed at all cost.

**Relation to previous predictive spiking models**. Previously, Boerlin et al. described how a population of spiking neurons can efficiently encode a dynamic input variable [10, 21, 22, 23]. In this work, we showed that a recurrent population of integrate-and-fire neurons with dynamics and connectivity tuned for efficient coding maintains a balance between excitation and inhibition, exhibits Poisson-like spike statistics [10], and is highly robust against perturbations such as neuronal cell death. However, we did not previously demonstrate a relation between efficient coding and neural oscillations. The main reason for this is that we always considered an idealised network, with instantaneous synapses. In this idealised network, only one cell fires at a time. As a result, oscillations are generally extremely weak and fast (with frequency equal to the population firing rate), and thus, completely washed out in a large network with added noise and/or heterogenous read-out weights. In contrast, in a network with finite synap delays, more than one neuron may fire per oscillation cycle, before the arrival of inhibition. As a result, oscillations are generally much stronger, even with significant added noise and heterogenous read-out weights (**Supp. fig. 4**).

**Attentional modulation**. Directing attention to a particular stimulus feature/location has been shown to increase the gamma-band synchronisation of visual neurons that respond to the attended feature/location [4]. **Fig. 8b-c** illustrates how such an effect could come about. Here, we show that attentional modulations that increase the strength of gamma-band oscillations will serve to increase perceptual discrimination of low contrast stimuli. Such attentional modulation could be achieved in a number of different ways, for example, by decreasing noise fluctuations, or modulating the effective gain of feed-forward or recurrent connections [24, 25].

In general, the way in which attention should modulate the network dynamics will depend on the stimulus statistics and task-setup. Future work, that considered higher dimensional sensory inputs (as well as competing ‘distractor’ stimuli), could allow us to investigate this question further.

**The benefits of noise**. With low noise, neural membrane potentials are highly correlated (**Fig. 4a**), and inhibition is not able to prevent multiple neurons firing together. To avoid this, we needed to add noise to neural membrane potentials (while simultaneously increasing spiking thresholds). With the right level of noise, fewer neurons fired on each oscillation cycle, resulting in increased coding performance. Too much noise, however, led to inconsistent information being encoded by different neurons, decreasing coding performance (**Fig. 4c**). This phenomena, where noise fluctuations increase the signal processing performance of a system, is often referred to as ‘stochastic resonance’ [26, 27], and has been observed in multiple sensory systems, including cat visual neurons [28, 29]. Previously however, stochastic resonance has usually been seen as a method to amplify sub-threshold sensory signals that would not normally drive neurons to spike. Here, in contrast, noise desynchronises neurons that receive similar recurrent inputs, increasing the coding efficiency of the population.

**Alternative functional roles for oscillations**. Neural oscillations have been hypothesised to fulfill a number of different functional roles, including feature binding [30], gating communication between different neural assemblies [31, 32, 33], encoding feed-forward and feed-back prediction errors [34, 35, 36] and facilitating ‘phase codes’ in which information is communicated via the timing of spikes relative to the ongoing oscillation cycle [37].

Many of these theories propose new ways in which oscillations encode incoming sensory information. In contrast, in our work network oscillations do not directly code for anything, but rather, are predicted as a consequence of efficient rate coding, an idea whose origins go back more than 50 years [8].

## Materials and Methods

**Efficient spiking network**. We consider a dynamical variable that evolves in time according to:

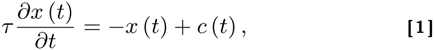

where *c*(*t*) is a time-varying external input or command variable, and *τ* is a fixed time constant. Our goal is to build a network of *N* neurons that take *c*(*t*) as input, and reproduce the trajectory of *x*(*t*). Specifically, we want to be able to read an estimate 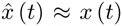 of the dynamical variable from the network’s spike trains *o*(*t*) = (*o*_1_(*t*), *o*_2_(*t*),…, *o_N_*(*t*)). These output spike trains are given by 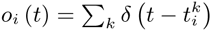, where 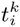 is the time of the *k^th^* spike in neuron *i*.

We first assume that the estimate, 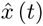, can be read out by a weighted leaky integration of spike trains:

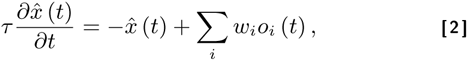

where *w_i_* is a constant read-out weight associated with the *i^th^* neuron.

We next assume that the network minimises the distance between *x*(*t*) and 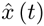 by optimising over spike times 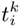. The network minimises the loss function,

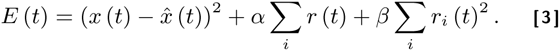

The first term in the loss function is the squared distance between the *x*(*t*) and 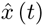. The second term and third term represent L1 and L2 penalties on the firing rate, respectively. *α* and *β* are constants that determine the size of the penalty. The time varying firing rate of the *i^th^* neuron is defined to by the differential equation:

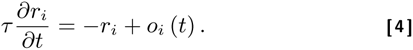

A neuron fires a spike at time *t* if it can reduce the instantaneous error *E*(*t*) (i.e. when *E*(*t***|**neuron *i* spikes) < *E*(*t***|**neuron *i* doesn’t spike)). This results in a spiking rule:

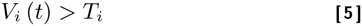

where,

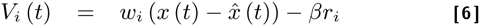

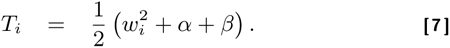

Since *V_i_*(*t*) is a time-varying variable, whereas *T_i_* is a constant, we identify the former with the *i^th^* neuron’s membrane potential *V_i_*(*t*), and the latter with its firing threshold *T_i_*.

To obtain the network dynamics, we take the derivative of each neuron’s membrane potential to obtain:

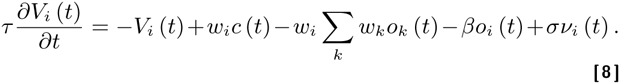

where *ν_i_*(*t*) corresponds to a white ‘background noise’, with unit variance (added for biological realism). Thus, the resultant dynamics are equivalent to a recurrent network of leaky integrate-and-fire (LIF) neurons, with leak, −*V* (*t*), feed-forward input, *w_i_c*(*t*), recurrent input, 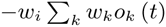, and self-inhibition (or reset), −*βo_i_*(*t*).

**Balanced network of inhibitory and excitatory neurons**. To construct a network that respects Dale’s law, we introduce a population of inhibitory neurons, that tracks the estimate encoded by the excitatory neurons, and provides recurrent feedback. For simplicity, we consider a network in which all read-out weights are positive. In our framework, this results in a particularly simple network architecture, in which a single population of excitatory neurons is recurrently connected to a population of inhibitory neurons (Figure 3a). For further discussion of different network architectures, see [10].

We first introduce a population of inhibitory neurons, that receive input from excitatory cells. The objective of the inhibitory population is to minimise the squared distance between excitatory and inhibitory neural reconstructions (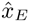, and 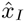, respectively), by optimising over spike times 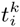. Thus, an inhibitory neuron spikes when it can reduce the loss function:

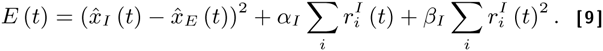

Following the same prescription as before, we obtain the following dynamics for the inhibitory neurons:

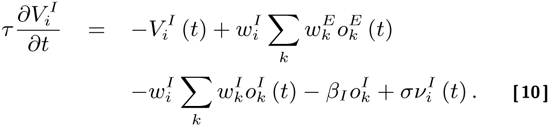

Thus, inhibitory neurons receive input from excitatory neurons (2nd term), and recurrent inhibition from other inhibitory neurons (3rd term).

Now, as the inhibitory reconstruction tracks the excitatory reconstruction, we can replace 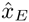 with 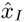, in the excitatory loss function, to obtain:

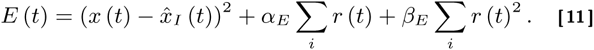

Following the same prescription once again, we obtain the following dynamics for the excitatory neurons:

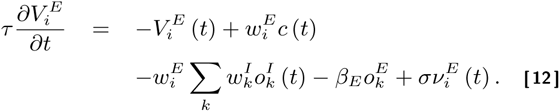

Thus, excitatory neurons receive excitatory feed-forward input (2nd term) and recurrent inhibitory input (3rd term).

**Synaptic dynamics**. To account for transmission delays and continuous synaptic dynamics we assume that each spike generates a continuous current input to other neurons, with dynamics described by the synaptic waveform, 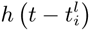. The shape of this waveform is given by:

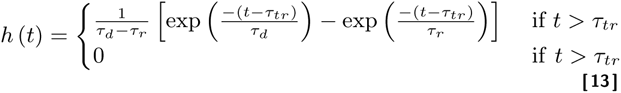

where *τ_r_* is the synaptic rise time, *τ_d_* is the decay time and *τ_tr_* is the transmission delay. The normalisation constant, *τ_d_* − *τ_r_*, ensures that 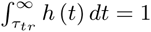. This profile is plotted in figure 3b, with *τ_r_* = 1ms, *τ_d_* = 3ms and *τ_tr_* = 1ms.

To incorporate continuous synaptic currents into the model, we alter equations 10 and 12, by replacing each of the recurrent spiking inputs (*o_k_* (*t*)) by the convolution of the spiking input and current waveform 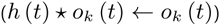.

**Simulation parameters**. For the simulations shown in **figures 2**, we considered a toy network of 3 neurons with equal read-out weights, *w_i_* = 1. The L1 spike cost was *α* = 0 and the L2 spike cost was set to *β* = 0.04. The read-out time constant was set to *τ* = 0.1*s*. The magnitude of injected membrane potential noise was set to *σ* = 0.02. In each case, network dynamics were computed from equation 8.

For the simulations shown in **figures 3-8**, we considered a larger network of 50 excitatory neurons, and 50 inhibitory neurons. All neurons had equal read-out weights, equal to *γ*_0_ = 1.2mV^1/2^. The L1 spike cost was set to 0. The L2 spike cost was set to *β* = 8.5mV. If we assume a spike threshold of −55mV, this corresponds to a resting potential of 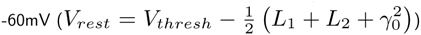, a reset of 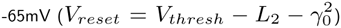, and post-synaptic potentials of 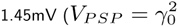; see [10] for details of scaling to biological parameters).

The magnitude of injected membrane potential noise was set to *σ* = 13’5mV. Network dynamics were computed from equations 10 and 12 (with the exception that recurrent inputs were convolved with the synaptic current waveform, *h* (*t*), described in equation 13).

**Algorithm**. Simulations were run using Runga-Kutta method, with discrete time steps of 0.5ms. For the ‘ideal’ network (i.e. with instantaneous synapses) only one neuron (with highest membrane potential) was allowed to fire a spike within each bin. Changing the temporal discretisation did not qualitatively alter our results.

**Stimulus details**. In **figure 2b**, the encoded variable, *x*, was obtained by low-pass filtering white noise with a 1st-order Butterworth filter, with cut-off frequency of 4Hz. After filtering, *x*, is rescaled to have mean of 3, and standard deviation of 1. In **figure 2c** the encoded variable is constant, *x* = 4. In **figure 3**, the encoded variable is obtained by low-pass filtering a white-noise input in the same way as for figure 2b, this time with a cut-off frequency of 2Hz. After filtering, *x*, is rescaled to have non-zero mean of 50, and standard deviation of 10. In **figure 4**, the ‘low’, ‘medium’ and ‘high’ amplitude inputs are *x* = 36, 48, and 60, respectively. In **figure 5** the encoded variable, *x*, is initially equal to 10, before increasing to a constant value of 30, 40, or 50 (blue, red and green plots respectively). In **figure 5b-d** and **figures 6-7**, the encoded variable was held constant at *x* = 50.

**Poisson model**. In **figure 3d-e**, we compare the efficient coding model to a rate model, in which neural firing rates vary as a function of the feed-forward input, *c*(*t*). To obtain firing rates, we computed the average firing rate of neurons in the recurrent model, for different values of the feed-forward input. This gave us empirical firing rates: *r* = *f*(*c*). Spiking responses were obtained by drawing from a Poisson distribution, with this firing rate. Later, we multiplied the firing rates by a constant factor, such that the mean-squared error was the same for the Poisson model and the efficient coding model.

**Varying the noise**. For the simulations shown in **figure 4**, we multiplied varied the injected membrane potential noise for all neurons by a constant factor. In general, varying the noise amplitude changes neural firing rates, leading to systematic estimation biases. To compensate for this, we adjusted the L2 spike cost for the inhibitory and excitatory neurons, so as to maintain zero estimation bias (or equivalently, to keep firing rates constant). For each noise level, we ran an initial simulation, modifying excitatory and inhibitory costs, *β_E_* and *β_I_*, in real time (via a stochastic gradient descent algorithm), until both the excitatory and inhibitory estimation biases converged to zero.

**Population firing rates**. To plot the population firing rate (**Fig. 5a & b**), we low-pass filtered neural spike trains using a first order Butterworth filter (with cut-off frequency of 5.5Hz, and 66Hz, for panels b and d, respectively), before averaging over neurons.

**Spectral analysis**. The spectrogram of the population firing rate, shown in **figure 5a** (lower panel), was computed using a short-time Fourier-transform, with a Hamming time window of 60ms (Matlab’s ‘spectrogram function’), Finally, the instantaneous power spectrum was low-pass filtered with a first-order Butterworth filter, with cut-off frequency 3Hz, The power spectrum of the population firing rate and neural membrane potentials was computed using the multi-taper method (using Matlab’s ‘pmtm’ function), with bandwidth chosen empirically to achieve a spectrum that varied smoothly with frequency.

**Excitatory and inhibitory currents**. To plot the currents shown in **Fig. 7b**, we divided the total excitatory and inhibitory input to each cell by a presumed membrane resistance of *R_m_* = 5MΩ (changing this value rescales the y-axis), We then defined peaks in excitatory and inhibitory currents as local maxima in the currents, separated by a drop an 80% drop in the current magnitude, Further, we only included peaks in inhibitory and excitatory currents that occurred within 15ms of each other.

**Discrimination threshold**. For the simulation shown in **figure 8**, we considered the performance of the network in discriminating between two 0,1s long stimulus segments, with equally spaced around the ‘low’ amplitude input (*x* = 48), From signal detection theory, a subject’s probability of selecting between two stimuli is given by: 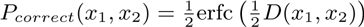, where erfc(*x*) is the cumulative error function, and *D*(*x*_1_, *x*_2_) is the normalized distance (or d-prime) between the distribution of estimates: 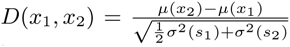, These quantities can be directly computed from the network output.

## ACKNOWLEDGMENTS

Support from the Basic Research Program of the National Research University Higher School of Economics is gratefully acknowledged by BSG.

